# Uneven Data Quality and the Earliest Occupation of Europe – The Case of Untermassfeld (Germany)

**DOI:** 10.1101/211268

**Authors:** Wil Roebroeks, Sabine Gaudzinski-Windheuser, Michael Baales, Ralf-Dietrich Kahlke

## Abstract

The database regarding the earliest occupation of Europe has increased significantly in quantity and quality of data points over the last two decades, mainly through the addition of new sites as a result of long-term systematic excavations and large-scale prospections of Early and early Middle Pleistocene exposures. The site distribution pattern suggests an ephemeral presence of hominins in the south of Europe from around one million years ago, with occasional short northward expansions along the western coastal areas when temperate conditions permitted. From around 600,000-700,000 years ago Acheulean artefacts appear in Europe and somewhat later hominin presence seems to pick up, with more sites and now some also present in colder climatic settings. It is again only later, around 350,000 years ago, that the first sites show up in more continental, central parts of Europe, east of the Rhine. A series of recent papers on the Early Pleistocene palaeontological site of Untermassfeld (Germany) makes claims that are of great interest for studies of earliest Europe and are at odds with the described pattern: the papers suggest that Untermassfeld has yielded stone tools and humanly modified faunal remains, evidence for a one million years old hominin presence in European continental mid-latitudes, and additional evidence that hominins were well-established in Europe already around that time period. Here we evaluate these claims and demonstrate that these studies are severely flawed in terms of data on provenance of the materials studied and in the interpretation of faunal remains and lithics as testifying to a hominin presence at the site. In actual fact any reference to the Untermassfeld site as an archaeological one is unwarranted. Furthermore, it is not the only European Early Pleistocene site where inferred evidence for hominin presence is problematic. The strength of the spatiotemporal patterns of hominin presence and absence depend on the quality of the data points we work with, and data base maintenance, including critical evaluation of new sites, is crucial to advance our knowledge of the expansions and contractions of hominin ranges during the Pleistocene.

## Introduction

### The earliest occupation of Europe

At the end of the Pleistocene human populations had colonized almost all corners of the world, in a process of range expansion from Africa which had started minimally by 1,8 million years ago. The spatiotemporal patterns of the traces of hominin presence and absence can inform on hominin adaptability and the dynamics of range expansions and contractions in the face of environmental changes of the Pleistocene. Europe has the best documented record in this regard, as a result of its long and intensive research history. That record is the more interesting as this *cul de sac* of the Eurasian landmass is large enough to display ecological differences from south to north and from west to east, which may have formed ecological clines and barriers during range expansion. Furthermore a rich database exists on Pleistocene climatic and environmental variability in this part of the hominin range which allows interpretation of patterns of hominin colonization against the background of fine-grained environmental changes [1–6].

Our database regarding the earliest occupation of Europe has increased significantly in quantity and quality of data points over the last two decades, mainly through the addition of new sites as a result of long-term systematic excavations (e.g. at Atapuerca, Spain [7, 8], with late Early Pleistocene finds, dating to more than 800 ka (1 ka = 1,000 years before present) and large-scale prospections of Early and early Middle Pleistocene exposures. A good example of such prospection work is the case of the Cromer Forest Bed formation in East Anglia, UK [9, 10], which yielded evidence for surprisingly early (up to c. 800 ka) - and possibly short-term - range expansions into northwestern Europe, up to 53 degrees North.

The hominin occupation of northwestern Europe before ~500-600 ka was on current evidence [3, 5, 11] highly sporadic, and probably the result of temporary range expansions of southern (circum-Mediterranean) populations when conditions ameliorated and permitted short northward movements, probably along its western coastal margins [12, 13]. In general terms in the whole of Europe there still seems to be a threshold for longer-term hominin settlement at around 500-600 ka, with a marked increase in the number of sites (now not only during temperate climatic intervals, but also during colder and drier phases) and the sizes of the assemblages. It is also from around 600-700 ka onward that we observe the first presence of Acheulean tools in Europe [14], about a million years later than their first appearance in eastern Africa [15]. The first occupants of Europe seem to have done without handaxes, the earliest European assemblages only comprising simple stone flakes, cores and core-like tools, with a lack of standardized design and usually with limited modification only. This gives special importance to the study of the geological context of inferred early sites: rocks can fracture naturally and edges can be modified by natural processes in sediments such as cryoturbation, transport and volcanic activities, and a wide variety of such processes has been documented to mimic hominin modification and to produce “artefact-like” geofacts, [16–19] (see also below, Discussion). Interestingly, the emergence of the Acheulean signal in southern [20] as well as northwestern Europe from 600-700 ka [14, 21, 22] onward is in the same time range as the current estimate for the beginning of the Neandertal lineage [23].

In the second half of the Middle Pleistocene, from around 350 ka onward, archaeological sites are not anymore limited to the western (Atlantic) and southern (Mediterranean influenced) regions of Europe, but also show up in its more continental areas, east of the river Rhine catchment area [6, 12, 24]: a range expansion into more challenging environments that might be related to the development of new strategies for survival [25], including the introduction of fire as a fixed part of the hominin technological repertoire [13, 26, 27].

The authors of a recent study of faunal material attributed to the late Early Pleistocene palaeontological site of Untermassfeld (Germany) [28] make two claims that are of great relevance in the context of the study of the earliest occupation of Europe. Firstly, they reiterate their previous [29–31] assertion that the Untermassfeld site has also yielded lithic artefacts and that hence Central Europe saw already a hominin presence about one million years ago. Secondly, they also claim to have identified in the faunal remains from the site numerous evidence of hominin butchering activities such as cut marks and intentional hammer stone-related bone breakage and identify exploitation patterns which suggest year round primary access to preferentially bison, horse, deer and megafauna species. The authors conclude that the stone tools and the humanly modified faunal remains from Untermassfeld provide evidence for the earliest hominin presence in European continental mid-latitudes at about 50 degrees North. Together with the evidence from inferred contemporaneous sites such as Vallparadis (Spain) [32] and Le Vallonet (France) [33] they see this as additional evidence that humans were well-established in Europe already one million years ago. If these claims are substantiated, the Untermassfeld site, a striking outlier in the geographical distribution of early Palaeolithic sites [19, 34] thus far and at odds with the spatio-temporal pattern described above, would indeed be a very interesting occurrence. The far-reaching implications of these claims, warrant an in-depth assessment of the arguments presented by these authors [28–31, 35].

The increase of the European archaeological record (see below: Discussion) has opened possibilities to study range expansions and contractions of hominin populations, including regional extinctions, during the Pleistocene [3, 36-39] and has allowed rephrasing the question “When was Europe first occupied?” into “How often was Europe abandoned after hominins first entered it?” [39]. Key to such studies is establishing the strength of the spatiotemporal patterns we work with. When studying the early occupation history of any given area, palaeoanthropologists’ first task is to critically assess the strength of the evidence for a presumed hominin presence, as well as to provide a solid framework for bracketing the age of such finds, i.e. lithics and fossils. The quality of these data determines the quality of the resulting spatiotemporal patterns of hominin presence and absence and hence the quality of the hypotheses developed to interpret these patterns. In a discipline in which the data about the exact provenance and geological context of finds is usually only obtainable through publications, researchers are banking heavily on the quality of fieldwork and of the resulting publications, including detailed description of provenance data and reproducibility of analyses carried out in the laboratory, which also entails accessibility of the finds at stake.

In this paper we will give a brief description of the Untermassfeld site, including its history of systematic excavations and analyses and its geological genesis. We will then evaluate the strength of the recent claims for a hominin presence at the site at the end of the Early Pleistocene in terms of lithics and faunal remains [28–31, 35] and end with a brief discussion of the current debate on the early hominin occupation of Europe.

### The Untermassfeld site: research history and geological setting

The Early Pleistocene fossil site of Untermassfeld, near the town of Meiningen in Southern Thuringia (Central Germany), was discovered in January 1978, and has since the discovery been the focus of continuous systematic fieldwork, carried out exclusively by the Senckenberg Research Station of Quaternary Palaeontology, Weimar (formerly the Institute of Quaternary Palaeontology, Weimar) (Fig 1). So far the field seasons include a total of 90 months of fieldwork (status as of January 1th, 2017). The three-dimensionally fine scale documented excavations (Fig 1) have produced a total of 17,882 catalogued specimens or specimen series including 14,224 fossil remains of large mammals from a total of 650 grid squares (1 m2 each; status as of January 1th, 2017), excavated to a maximum depth of 5.50 m below ground level. Depending on the find density and conservation status, fossils were taken out of the embedding sediment as individual pieces, removed in plaster jackets to ensure stability, or lifted by crane in large blocks from the excavation site. During the fieldwork free periods, the active sections of the site were covered with 1 tonne weighing steel plates measuring 2 x 4 m, to minimize unauthorized access by third parties.

**Figure 1.**
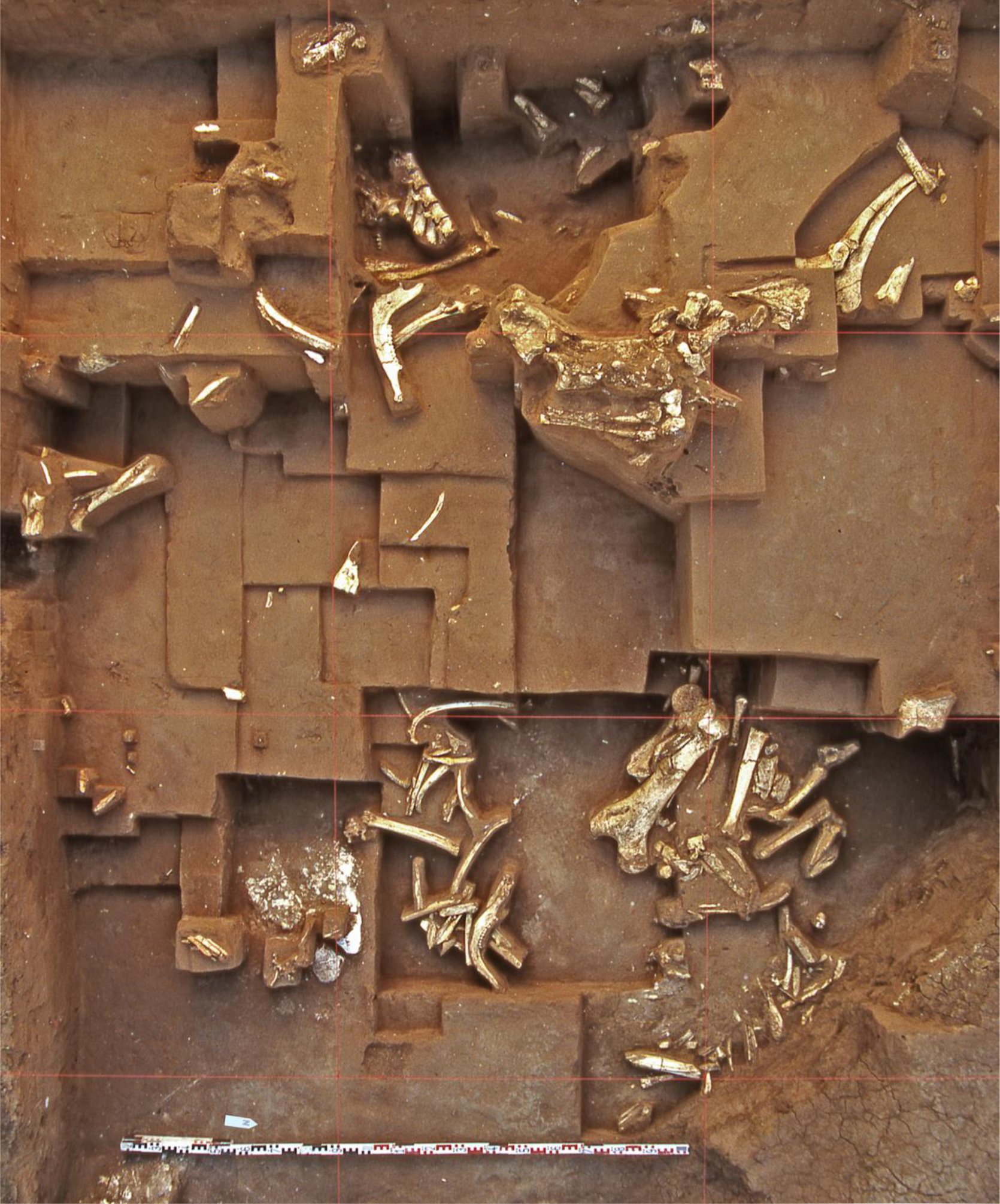
Untermassfeld research excavation (August 2002), excavation area with grid squares 21 - 24, 47 - 50, 154 - 158, 1.18 - 2.12 m below site 0-level, exposed finds of predominantly *Bison menneri, Dama nestii vallonnetensis, Eucladoceros giulii* and *Hippopotamus antiquus.*

The fossil assemblage recovered at Untermassfeld represents the most complete vertebrate record of the late Early Pleistocene in the Western Palaearctic, and includes numerous new discovered taxa. Importantly, this fauna has provided the base for establishing the Epivillafranchian as an independent biochron for the time period 1.2 to 0.9 Ma in the Western Palaearctic [6, 40, 41]. Untermassfeld is studied by an interdisciplinary research group, with 49 researchers from 31 institutions in 12 countries involved (status as of January 1th, 2017; www.senckenberg.de/root/index.php?page_id=2915. See [42–45] for detailed information on research history and the present results of this interdisciplinary research group). Not one of the authors of the various papers claiming a hominin presence at the site [28-31, 35] has ever been involved in the Untermassfeld project in whatever way.

At Untermassfeld the eastern slope of the Werra river valley cuts Triassic formations. Upstream of the fossiliferous site, in its immediate vicinity as well as in the subsoil, Middle Triassic calcareous rocks crop out (Fig 2). This so-called Muschelkalk formation occasionally contains thin layers of silicified greyish to greyish-brown chert. Triassic limestone and chert are frequently found in Early Pleistocene gravels cut by the fossiliferous layers of the site, as well as in all other Pleistocene fluviatile deposits of the middle Werra valley. The lithological characteristics of the sands of the Untermassfeld fossil site demonstrate that they are floodplain deposits, accumulated as a result of powerful high flood events [43, 46]. The deposition of the vertebrate and non-vertebrate faunal assemblage at the site occurred under the lee of an extended clastic mudflow fan that had poured from the adjacent valley slope into the river channel (Fig 2). Upstream of the excavation site a narrow part of the valley caused strong turbulences as well as a rapid rise of the upstream water level.

**Figure 2.**
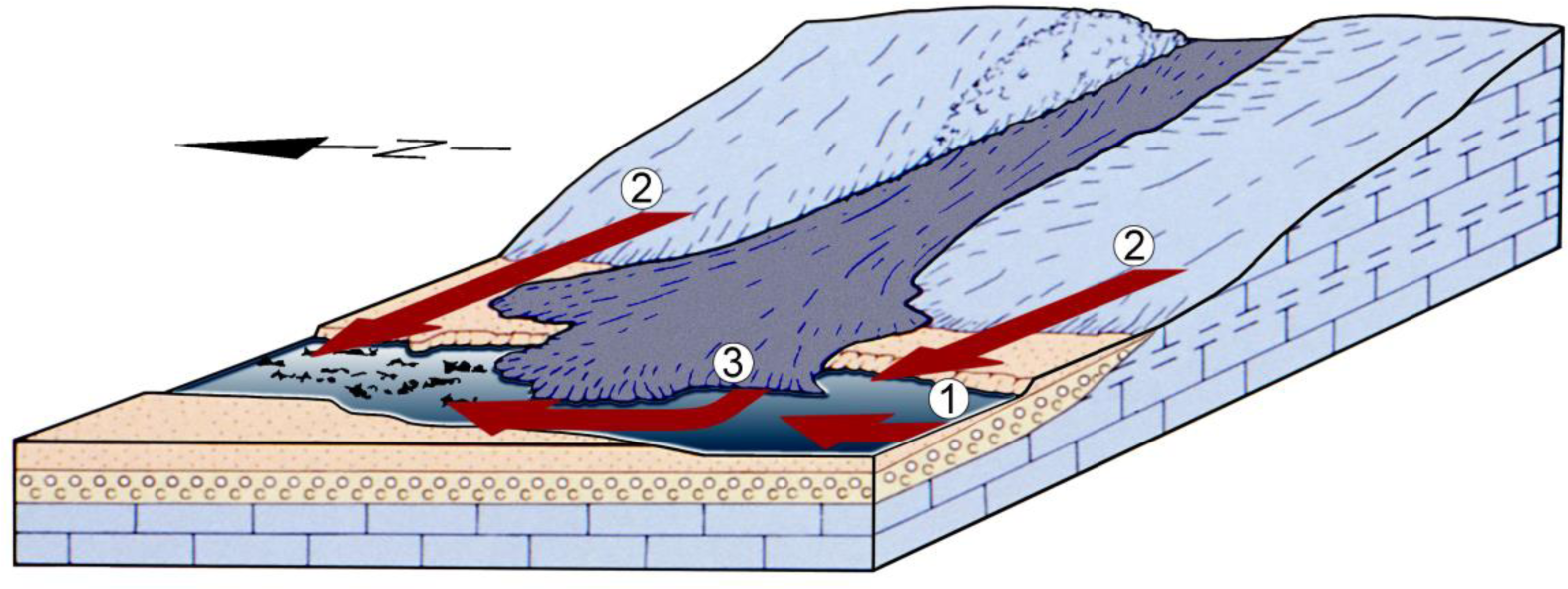
Schematic reconstruction of the Untermassfeld area during formation of the site: Geological sequence with Middle Triassic carbonates (brick symbol), fluviatile gravels (circles) and sands (dots), mass accumulation of carcass remains under the lee of an extended clastic mudflow fan (dashed). Arrows indicate the origin of limestone and chert rocks transported into the bone bearing sands: 1 - from the upper reaches of the Werra River and eroded subsoil of the river. 2 – from adjacent slopes of the valley. 3 – from the mudflow fan within the site itself (Drawing based on [43]).

During the formation of the find bearing deposits, reconstructed with data obtained during excavations and corresponding continuous geological surveys, rocks of Triassic limestone and chert were transported into the fossileferous deposits from various sources: the upper reaches of the river, the slopes of the valley, the eroded and reworked subsoil of the site and the clastic mudflow fan within the site itself (Fig 2). The re-deposition of Triassic material took place by means of high-energy processes that caused mechanical splintering and breakage of the corresponding rocks.

The Untermassfeld project has from its very beginning explicitly taken into consideration a possible presence of traces of hominin activities [42, 43] and lithics were continuously inspected for traces of hominin modification ([46], Plate 12,2), as were the faunal remains from the site (the first director of the excavations at the site, H.-D. Kahlke, was trained as an archaeologist and had worked at many Palaeolithic sites, including Zhoukoudian in China [47]). However, as repeatedly emphasized in publications on the site, no traces of a hominin presence whatsoever could be identified thus far [43, 48]. How then, does the material presented by Landeck and colleagues [28–31, 35] fit in the Untermassfeld evidence?

## Materials and methods

In order to evaluate the recent claims regarding a hominin presence at the sites we studied the material published in the articles at stake, as far as that was available: a small sample of faunal remains and eleven lithic objects. (why the examined sample is so small will be explained below). Zooarchaeological analyses of the faunal material focussed on the identification of surface modifications: in order to evaluate the claims made by Landeck and Garcia Garriga the identification of hominin induced modifications in the form of cut-marks and traces of marrow extraction was the focus of our attention. Bone surfaces were studied using a Leica reflected-light microscope with a magnification of up to 32x. All traces were registered per bone and recorded by anatomical position. Diagnostic criteria were used to identify hominin induced cut-marks [49, 50] and anthropogenic fractures [51–53]. M.B. and W.R. studied the lithics, in order to assess the artificial character of the finds, using hand lenses with magnification up to 12x. Besides basic measurements, types of raw material, character and numbers of negative scars, presence/absence of striking platforms, and cortical presence were studied.

To explain why the available sample of faunal remains and lithic finds is so small we need to briefly discuss the provenance of the material published by Landeck, Garcia Garriga and colleagues as well as the information on where it is deposited for other researchers to access. The sample presented by the referred authors [28–31, 35] allegedly consists of a number of 419 faunal remains and 256 stone objects. The provenance of the overwhelming bulk of the faunal and lithic materials however is unclear, as at no time in the site’s long excavation history any other regular excavations or other types of find recovery occurred. A recent paper in the *Journal of Human Evolution* describes faunal remains said to come from the Untermassfeld site [28], even though its authors were never involved in the excavations or in the investigations of the Untermassfeld material. Photos of the excavation which appear in their publication [28] were taken by a "tourist" (their Fig. 1) and by unknown persons who illegally gained access to the site (their Fig. 2, right). Regarding the origin of the published finds [28], the authors claim to have studied Untermassfeld faunal remains present in a “Schleusingen collection” (2016,58), assembled during "archaeological rescue operations at the Untermassfeld site in the late 70s and early 80s”. However, no such operations ever occurred at the site, and the Schleusingen collection either does not exist or is only known, and accessible, to Landeck and Garcia Garriga. Indeed, even the claim that the Untermassfeld site is the actual source of all the bone fragments is questionable. However, one fragment published by Landeck and Garcia Garriga (2016, Fig. 6c), the distal portion of a right metacarpal bone of *Dama nestii vallonnetensis,* does originate from the Untermassfeld excavation site: it was stolen from the site between 12:00 hrs on the 29th of May and 12:30 hrs on the 2nd of June, 2009, i.e. during the Pentecost 2009 weekend. It was broken when forcibly removed from a bone concentration (square Q 30; Fig 3a). Fragments of the proximal bone portion remaining in the layer (Fig 3b) were later recovered during official (licensed and documented) excavation work and then glued, preserved and catalogued [Senckenberg Research Station of Quaternary Palaeontology, Weimar, Untermassfeld Collection IQW 2010/ 30 740 (Mei 29 902)]. Both the theft and vandalism were reported to the responsible police authority (case number KPI Suhl / 1707-008127-09/1) and corresponding investigations were carried out by the Meiningen public prosecution office (case number 560 UJs 11134/09 60). Thefts and illegal excavation activities at the site had to be reported to the police also in the years 2002, 2006, 2007, 2009, 2010, 2011, and 2012.

**Figure 3.**
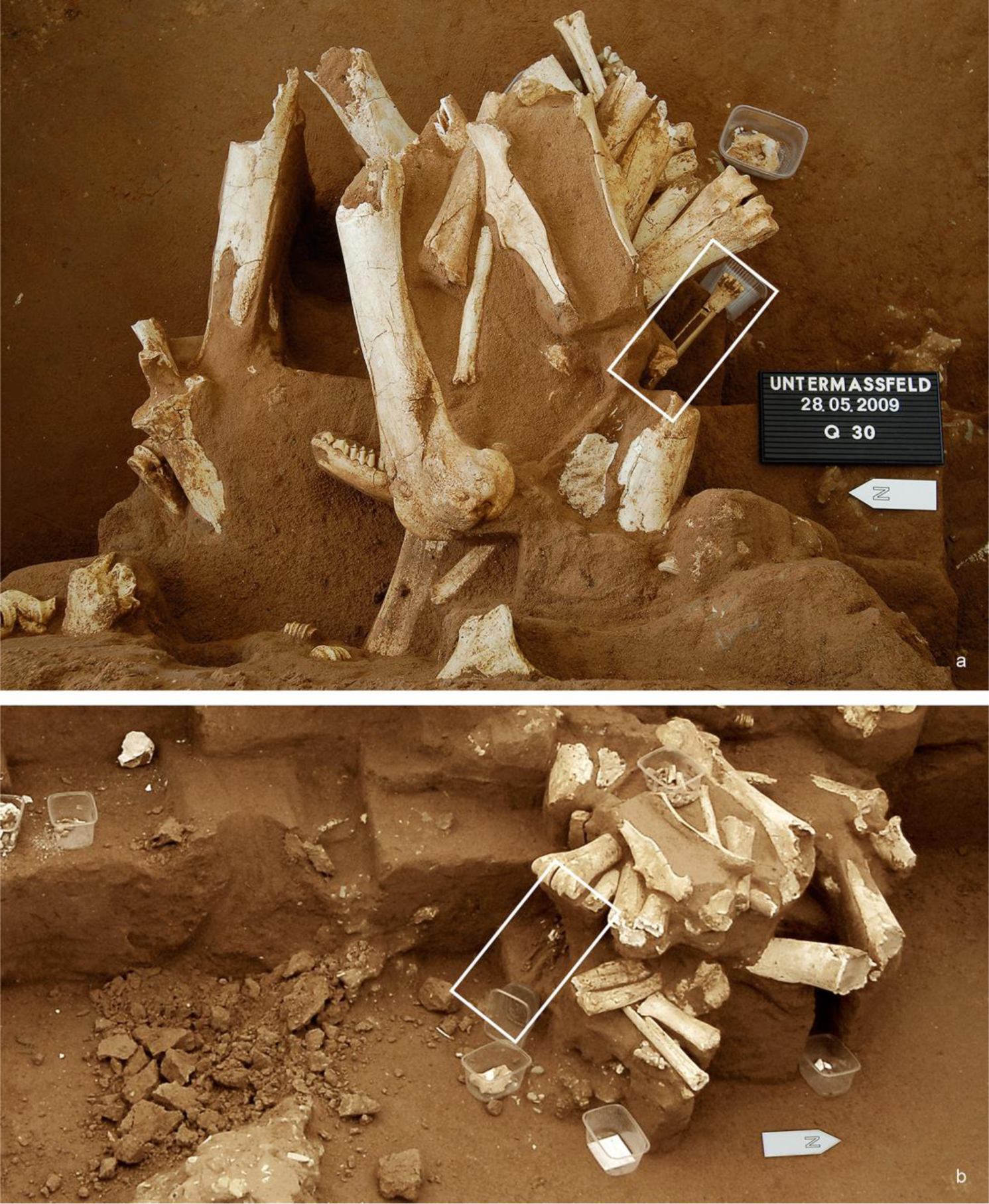
Untermassfeld research excavation covered by excavation tent: a – Undamaged polyspecific bone concentration exposed in square 30 (seen from above) with complete right metacarpal bone (rectangle) of a medium sized deer (*Dama nestii vallonnetensis*), 28^th^ of May, 2009. b – Damaged excavation view (picture of the same concentration but taken from different angle), distal part of the deer metapodial broken off, splintered proximal part still within the sediment block (rectangle), 2^nd^ of June, 2009.

Five years later, in March 2014, two packages were anonymously delivered to the Natural History Museum Schloss Bertholdsburg, Schleusingen (Thuringia). Along with an accompanying anonymous letter, the packages contained bone and rock material allegedly originating from the Untermassfeld site (personal communication by R. Werneburg, director of the Schleusingen museum, to R.-D. K. on the 22nd of July, 2014). Information on the circumstances surrounding the recovery of these specimens or their find locations was not included. One of the packages contained the above-mentioned metapodial fragment from *Dama nestii vallonnetensis*. This fragment, according to Landeck and Garcia Garriga [28] part of an assemblage recovered in the late 1970s and early 1980s, was still in the ground decades later, and stolen as an incomplete fragment from a 2009 excavation area (Fig 3). Fig 4 shows the regained distal metapodial fragment refitted to the proximal portion of the same skeletal element, recovered by the Senckenberg excavators. The whereabouts of the freshly broken, but still missing, portions of the bone remain unknown. The packages delivered to the Natural History Museum Schloss Bertholdsburg were transferred to the Senckenberg Research Station at Weimar. In these packages, further bone fragments published by Landeck and Garcia Garriga (2016, Figs. 3.b, 4.a, 4.e, 5.a, 5.e, 6.d, 7.d, 7.g, 8.a) were identified, in total 10 out of the 36 illustrated in their *Journal of Human Evolution* paper.

**Figure 4.**
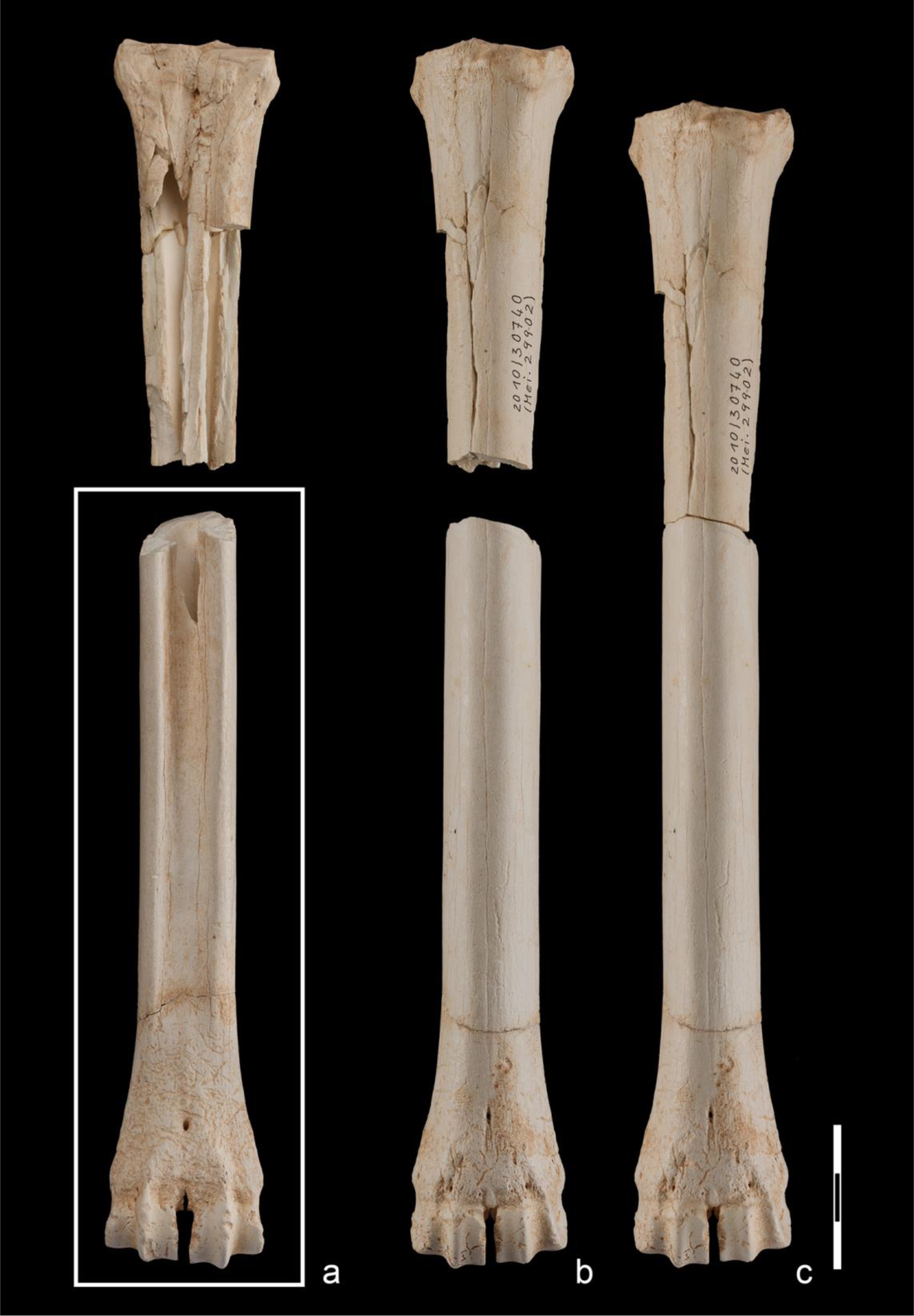
Untermassfeld, right metacarpal bone of *Dama nestii vallonntensis*: a-b - Proximal part of the bone: damaged during illegal activities, and then recovered during an official excavation, then prepared and cataloged. a-b - Distal part of the bone: broken off from an exposed polyspecific bone concentration in 2009 (rectangle; cf. Fig. 3a), published by Landeck and Garcia Garriga ([28], Fig. 6c), and anonymously transferred to the Natural History Museum Schleusingen. c – Excavated and stolen parts of the metapodial re-joined (now in the Untermassfeld collection of Senckenberg Weimar) (scale 3cm).

Given their facies, i.e. colour, degree of mineralization and brittleness, it is likely that these bones, like the deer metacarpal, also come from the Untermassfeld site. If we *assume* that the bone fragments in the packages (n=64) were part of the assemblage studied by Landeck and Garcia Garriga (2016: Table 1), the whereabouts of 355 pieces, i.e. 84% of the material allegedly studied, is presently unknown.

The provenance as well as the whereabouts of the majority of the published lithics are totally unclear: Garcia Garriga et al.[30] claim to have studied 256 lithic objects from Untermassfeld, without any documentation of where and how these were obtained. An online document from 2008 [35] simply states that “Nearly all of the lithic implements are very small and apparently have been overlooked by the excavating palaeontologists“. In a published exchange between Baales [54] and Landeck and Garcia Garriga [31] regarding the unclear provenance of the rock fragments Landeck and Garcia simply state that the lithics which they interpret as artefacts “…were recovered *in situ* from the Early Pleistocene sediments …which also included the large-mammal remains” [31] - without any information on the location of the finds and without any form of description of a relevant section whatsoever. It is also unclear where this material is deposited. Landeck [35] states that “Objects are handed over to the excavator Dr. R.D. Kahlke (Forschungsstation für Quartärpaläontologie, Senckenberg-Institut Weimar) and are stored at the Thüringer Landesamt für Denkmalpflege (Weimar)“. According to a later publication [29], a „small selection of specimens is preserved at the Thüringer Landesamt für Denkmalpflege (Weimar)“. (2010:1230). The Thüringer Landesamt für Denkmalpflege however does not hold lithics from Untermassfeld ( pers comm email Dr T. Schüler, Landesamt für Denkmalpflege und Archäologie, Weimar, September 6th 2016) and no lithics were handed over to the Senckenberg Research Station of Quaternary Palaeontology. However, the two packages with the faunal remains delivered to the Schleusingen museum in 2014 contained eleven small rock fragments, all of which we identified as having been published as artefacts by Garcia Garriga et al. [30] (their Figs 5a, 5c-5e, 6a-6d, 7c-e).

In summary, it must be concluded that the information given by Landeck and Garcia Garriga [28, 30] about the origins of the published lithic and faunal specimens is dubious, incomplete and at in one case demonstrably false. Furthermore, the publications provide no and/or incorrect data on where the studied material is stored and hence where it can be accessed by other researchers. For our analysis the only material we had at our disposal was the material in the anonymously delivered packages, momentarily stored at the Senckenberg Research Station of Quaternary Palaeontology at Weimar.

## Results

### Faunal remains

Landeck and Garcia Garriga infer hominin presence at the site on the basis of the presence of cut marks and traces of intentional hammerstone-related bone breakage on the bones of large mammals. This is interpreted as indicative of regular primary access to the carcasses of large mammals, butchering of complete animals and exhaustive exploitation through marrow processing. The bone fragments indeed were found to display surface modifications, but none of these are convincing in terms of manipulation by Pleistocene hominins. The problems that are apparent with the identification of presumable cut-marks published by Landeck and Garcia Garriga are manifold. The authors for instance misidentify modern striae as ancient cut-marks, as illustrated by a femur shaft fragment of a medium sized mammal (Fig 5). The surface damage is characterized by blunt, undefined shoulders that merge into the surrounding bone surface. The modification starts as very shallow striation in a modern break and is interrupted by a modern planar removal of the bone surface. Landeck and Garcia Garriga [28] interpret this damage as a defleshing cut-mark, overlapped by a tooth-mark, indicating primary access to carcasses by hominins ([28], 58, 63, Fig. 3b). Carnivore damage (cf. [55]) is evident on the edges of the bone fragment (Fig 5b).

**Figure 5.**
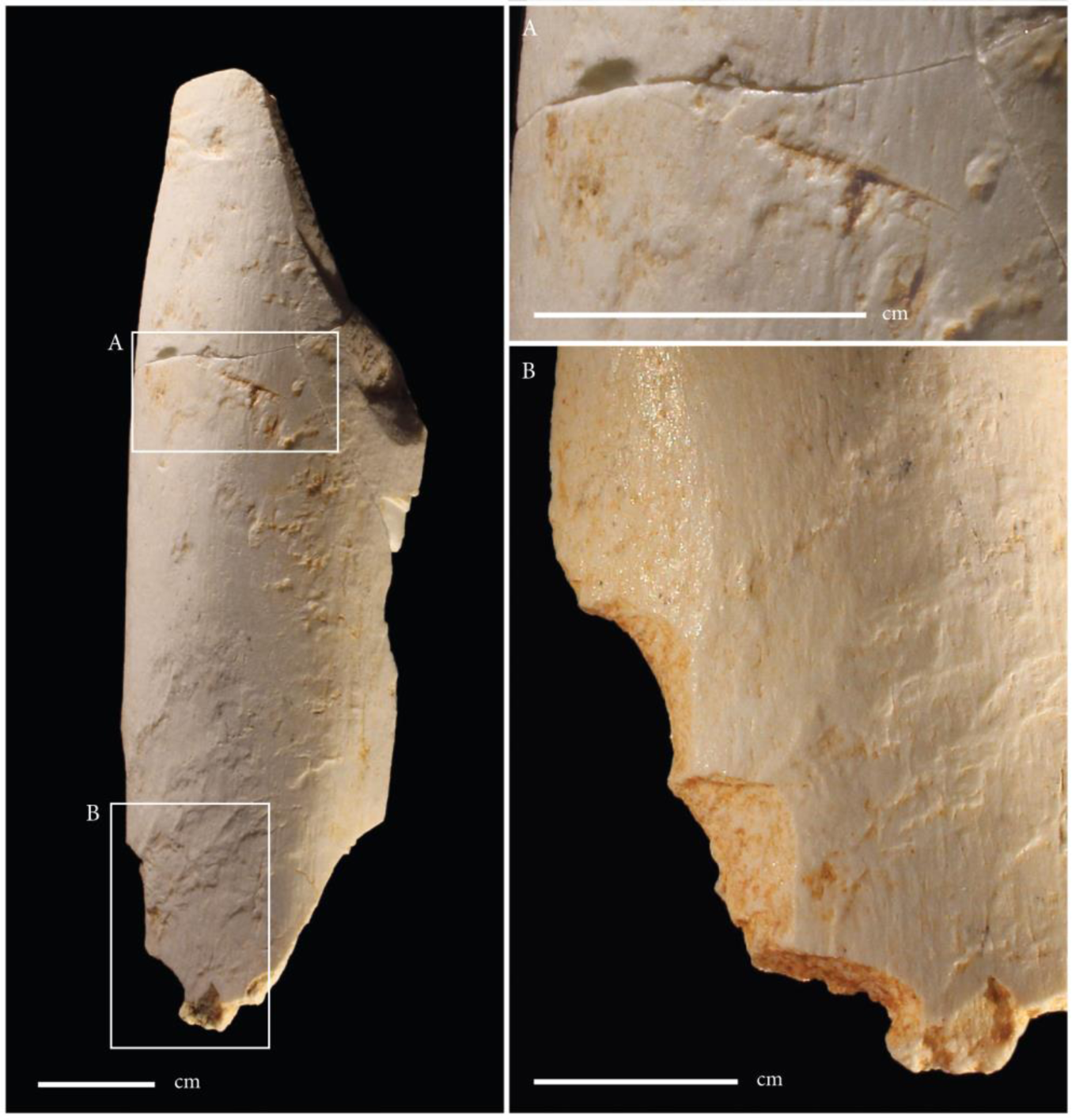
Femur fragment of a medium-sized mammal. The surface damage is characterized by blunt, undefined shoulders that merge into the surrounding bone surface. The modification starts as very shallow striation in a modern break and is interrupted by a modern planar removal of the bone surface. The lower left part of the modern planar surface removal is filled with red sediment, as well as the deeper section of the modification. Landeck and Garcia Garriga interpret this damage as a defleshing cut-mark, overlapped by a tooth-mark, indicating primary access to carcasses by hominins ([28] 58, 63, Fig. 3b). B. Carnivore damage is evident on the edges of the bone fragment.

A left calcaneus of *Stephanorhinus hundsheimensis* (Fig 6) with modern striae serves as a further example. Landeck and Garcia Garriga published these traces as hominin cut-marks providing evidence for the butchery of mega-fauna to obtain muscle meat ([28], 58, Fig. 4a). Moreover rodent modifications (cf were interpreted as being of anthropogenic origin (Fig 7) as exemplified by a right metacarpus of *Dama nestii vallonnetensis*: typical rodent modifications (cf. [56] Fig. 17) are present on the cranial face of the bone (Fig 7a), published as traces of skinning by Landeck and Garcia Garriga, ([28], 59, Fig. 6c). Additional striae, also published as being indicative of skinning, were located among an amalgam of shallow striae and surface modifications which result from root etching [57] (Fig 7b). The distal part of the bone also carries modifications caused by root etching and by rodent activities (Fig. 7c). In addition surface modifications found among an amalgam of traces of similar morphology were isolated in order to argue for the presence of cut marks (Fig 7b). In some cases, the published traces could not even be detected on the bones as in the case of a right astragalus of *Bison menneri* ([28], Fig 6d). Interpretations based on the anatomical location of inferred cut-marks are equally problematic. The interpretation of cut-marks on the posterior plantar tuber calcanei of the rhino *Stephanorhinus hundsheimensis* (Fig 6) ([28], 58, Fig. 4a) and on the diaphysis of a metapodial ([28], 58, Fig. 4e) as resulting from the exploitation of muscle meat is anatomically incorrect, since no muscle meat is present there nor can adjacent meat be detached by cutting at these locations. In brief, the claims made regarding the presence of faunal remains indicative of hominin butchering activities at Untermassfeld are unsubstantiated.

**Figure 6.**
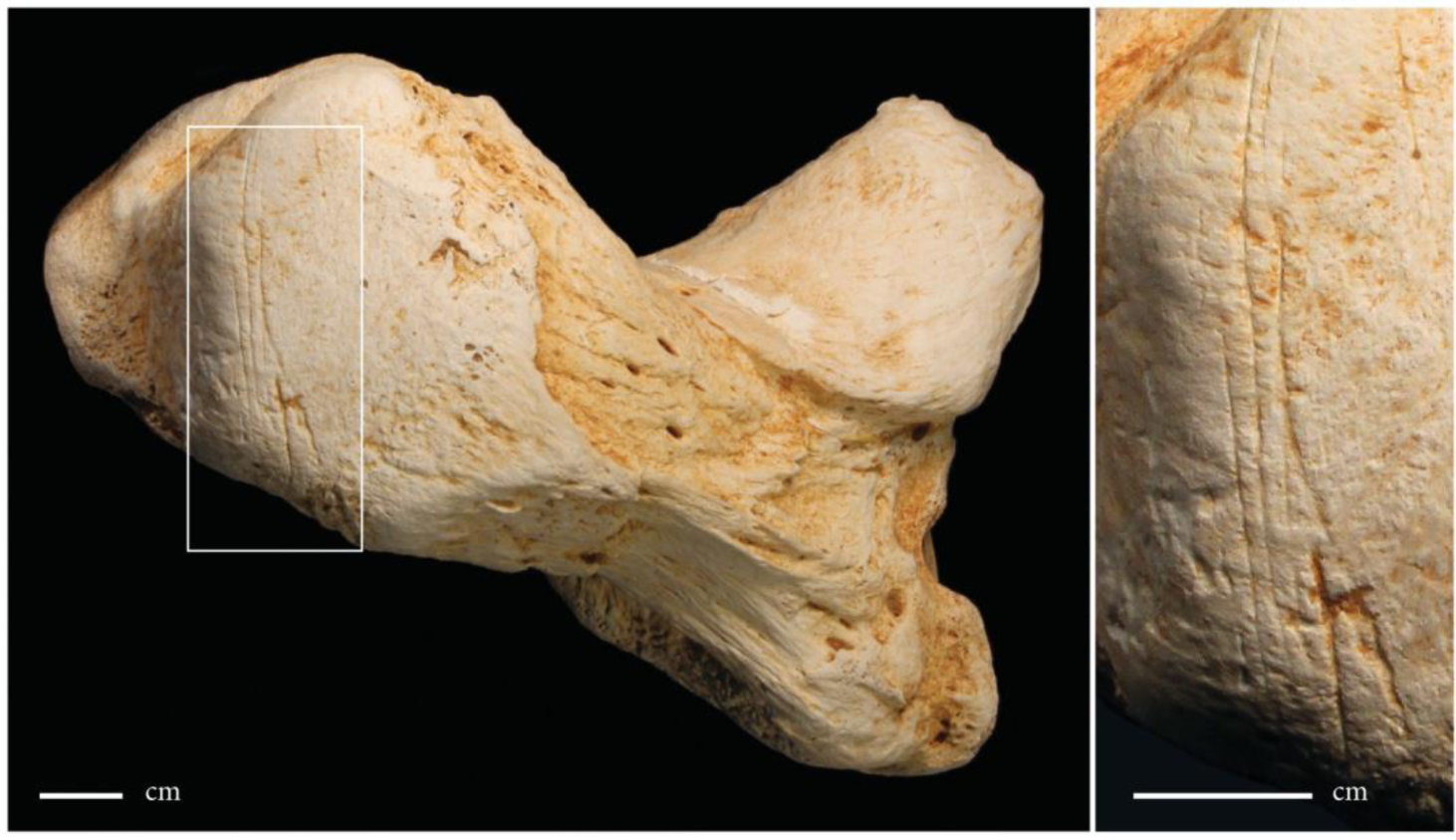
Left calcaneus of *Stephanorhinus hundsheimensis* with modern striae, partly filled with red sediment. Landeck and Garcia Garriga published these traces as hominin cut-marks providing evidence for the butchery of mega-fauna to obtain muscle meat ([28], 58, Fig. 4a).

**Figure 7.**
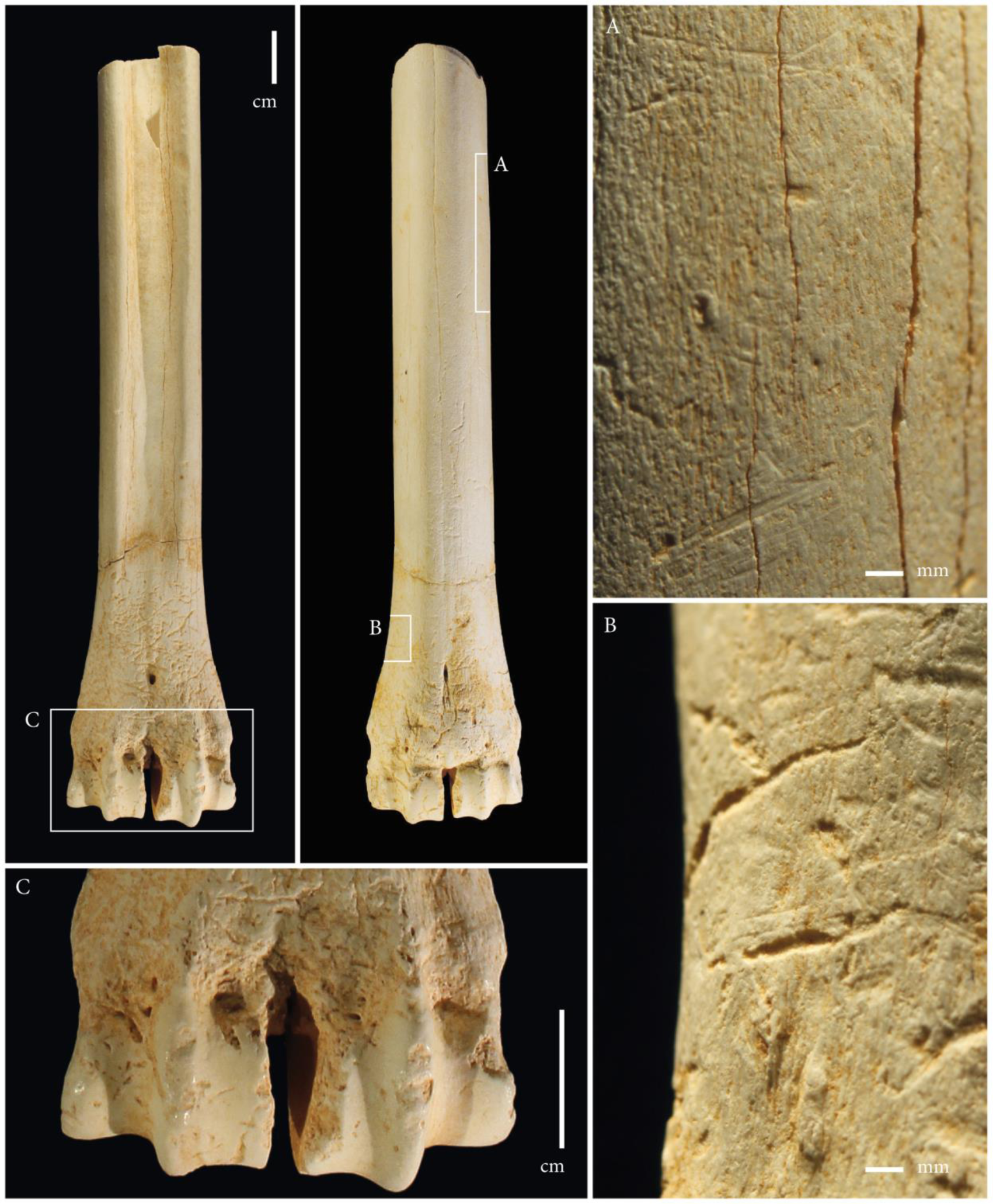
Right metacarpus of *Dama nestii vallonnetensis*. A. Typical rodent modifications could be observed on the cranial face of the bone, published as traces of skinning by Landeck and Garcia Garriga, ([28], 59, Fig. 6c). B. Additional striae, also published as being indicative of skinning, were located among an amalgam of shallow striae and surface modifications due to root etching. C. The distal part of the bone carries modifications due to root etching and rodents.

### Lithics

According to Landeck and Garcia Garriga, the inferred access to large mammal carcasses was facilitated through the use of “Mode 1” small microlithic tools (average length of retouched flakes 19,1 mm, of flakes 22,8 mm), which were produced in “short chaînes opératoires” using a systematised bipolar on an anvil knapping technique” and made on material “locally sourced from nearby streams and outcrops” ([30], 77-78). The unknown provenance and whereabouts of the small assemblage have already been addressed above. The packages with the faunal remains delivered to the Schleusingen museum also contained eleven small rock fragments, all published as artefacts by Garcia Garriga et al. [30] (their Figs 5a, 5c-5e, 6a-6d, 7c-e) and momentarily stored at the Senckenberg Research Station of Quaternary Palaeontology, Weimar. There are six chert (*Hornstein*) fragments and five limestone pieces, all sourcable to the local Triassic *Muschelkalk* deposits. To illustrate the character of these finds, we refer the reader to the original publication by Garcia Garriga et al. [30] with pictures of the objects, six of which we publish here too. Our Figures 8 to 10 show objects interpreted as cores “knapped on an anvil”: Fig 8 illustrates a broken chert piece, published as a “tabular chert core knapped on an anvil” ( [30] Fig 5e), our Fig 9 displays various views of a “pyramidal chert core”, with cortical remains (compare Fig. 5a in Garcia Garriga et al. 2013, only 22 mm in its largest dimension - and published with an incorrect scale bar), while Fig 10 shows another inferred core (compare Fig 5c in [30]). The objects illustrated in Figs 11 to 13 are interpreted by Garcia Garriga et al. [30] as flakes. Fig 11 shows a shattered limestone piece, with no bulbus, no platform, no retouch and with irregularly crushed edges, originally described as displaying “regular and continuous retouches at a steep angle” ([30], p.81; [30], Fig 7e). The lithic fragments displayed in Fig 12 and 13 (their [30]Figs 6b and 6d) have been published as chert flakes produced in a “bipolar on an anvil technique”. Not one of the eleven pieces has a clearly visible bulb of percussion, while a co-occurrence of a definable striking platform with scars of previous removals is also absent. The authors‘ own statement that part of the assemblage consists of „…pieces of shatter or chunks which lack characteristic fracture propagation characteristics“ [35] certainly applies to these eleven pieces, that in our view need to be interpreted as clear geofacts (cf. [19]). As detailed elsewhere [43, 46] and as mentioned above, the find bearing sands contain large amounts of Middle Triassic limestone debris in the form of chert and limestone fragments from the river valley slopes upstream and in the vicinity of the site, as well from the coarse clastic mudflow fan which moved downward from the slope into the fluvial deposits (Fig 2). These fragments are in the same size range and of the same facies and morphology as the “small artefacts” published by Garcia Garriga et al. [30] and are very abundant in the find bearing deposits. To illustrate this, our Fig 14 shows two boxes with such rock fragments retrieved during preparation of a sediment block of approximately 50x35x25 cm, containing the skull of a juvenile hyena, *Pachycrocutra brevirostris*, excavated in 1993, likewise illustrated in Fig 14. The figure underlines the abundant presence of shattered pieces of limestone and chert in the find bearing deposits (as well as the careful documentation, preparation and storage all finds from the Untermassfeld site).

The find location(s) of the lithic fragments published by Garcia Garriga and colleagues is undocumented and unknown, but the eleven pieces available perfectly fit within the natural background of shattered pieces from the *Muschelkalk* deposits both within and above the find bearing deposits, both in terms of their raw materials as well the natural modifications observable on these natural background objects (Fig 14). We conclude that the lithic pieces do not testify to any form of hominin modification, the lithic assemblage simply being the result of selecting naturally broken pieces from locally occurring deposits or from comparable adjacent outcrops in the Werra valley.

**Figure 8.**
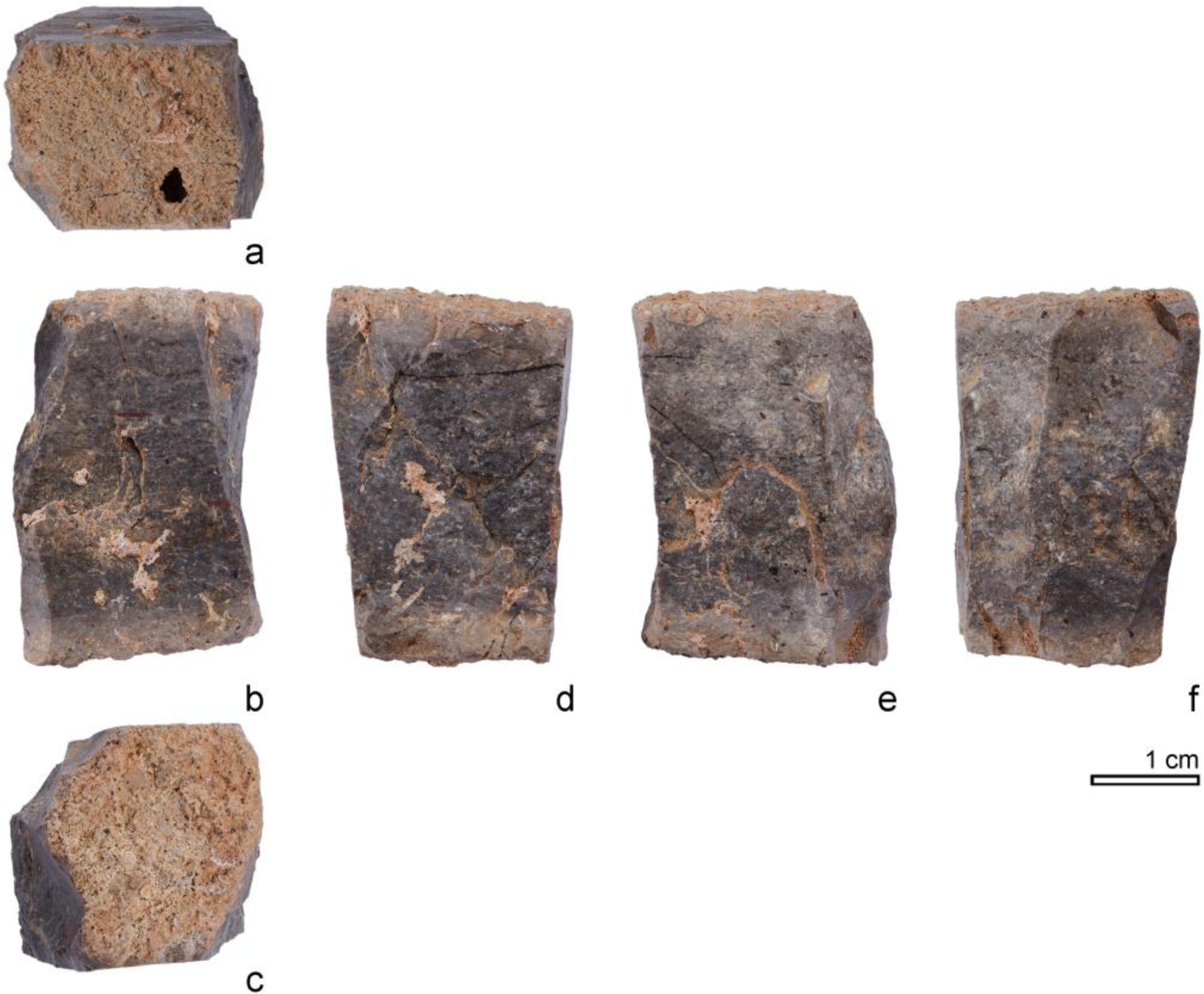
Chert fragment, published as a “tabular chert core knapped on an anvil” ([30], Fig 5e).

**Figure 9.**
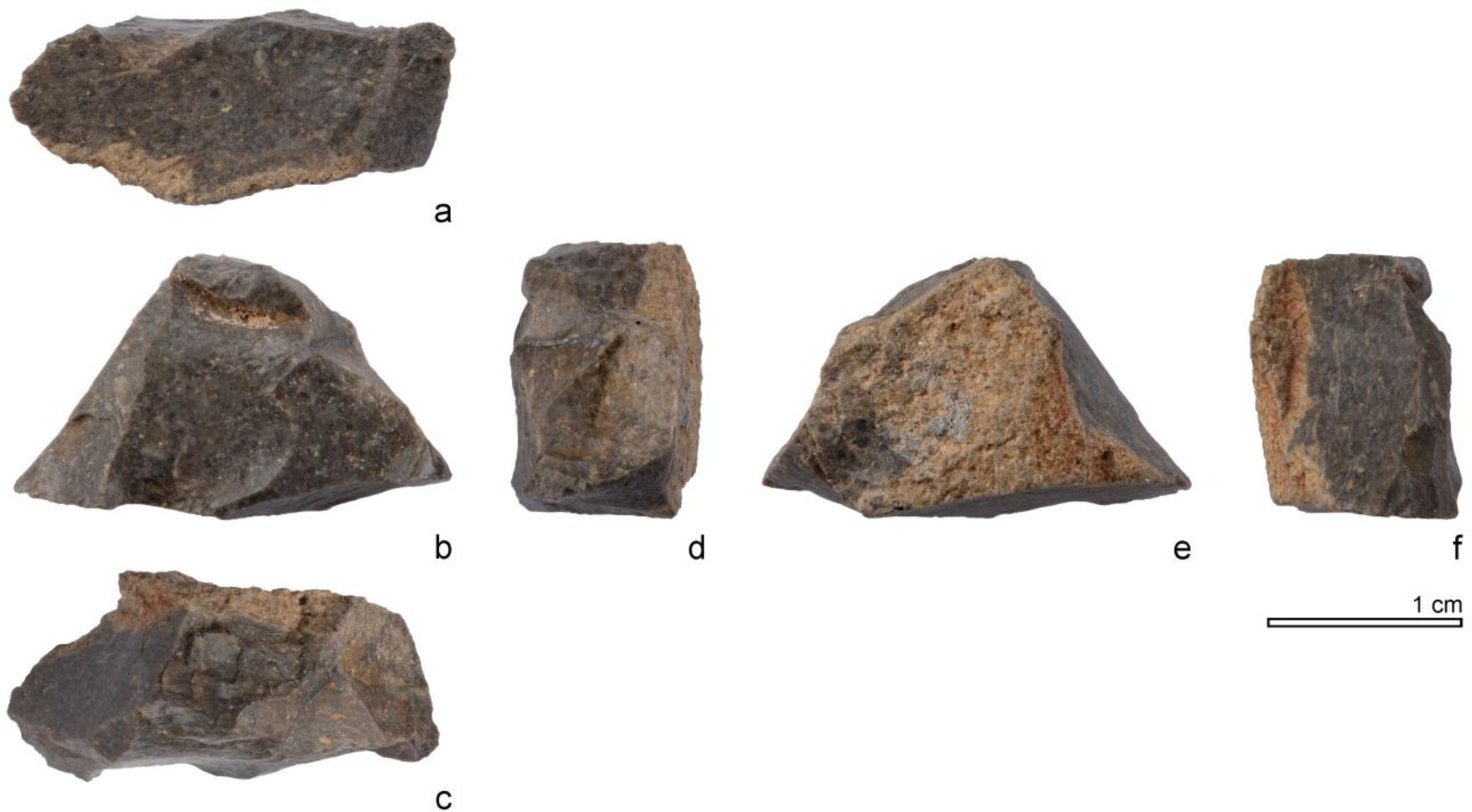
Chert fragment published as a “pyramidal chert core” with cortical remains (compare Fig. 5a in [30]), only 22 mm in its largest dimension.

**Figure 10.**
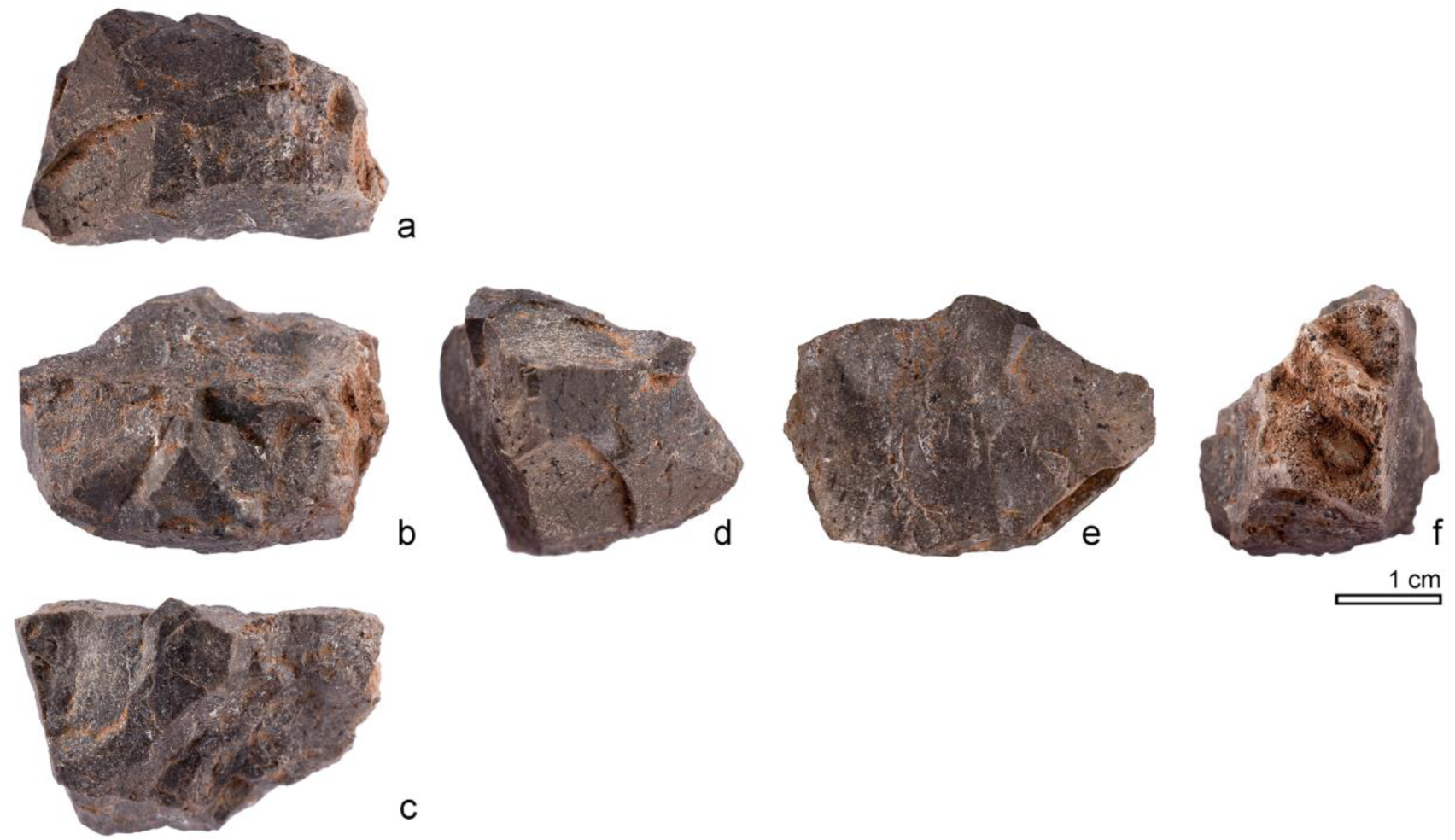
Silicified limestone fragment interpreted as a core ”knapped on an anvil” (compare Fig 5c in [30]).

**Figure 11.**
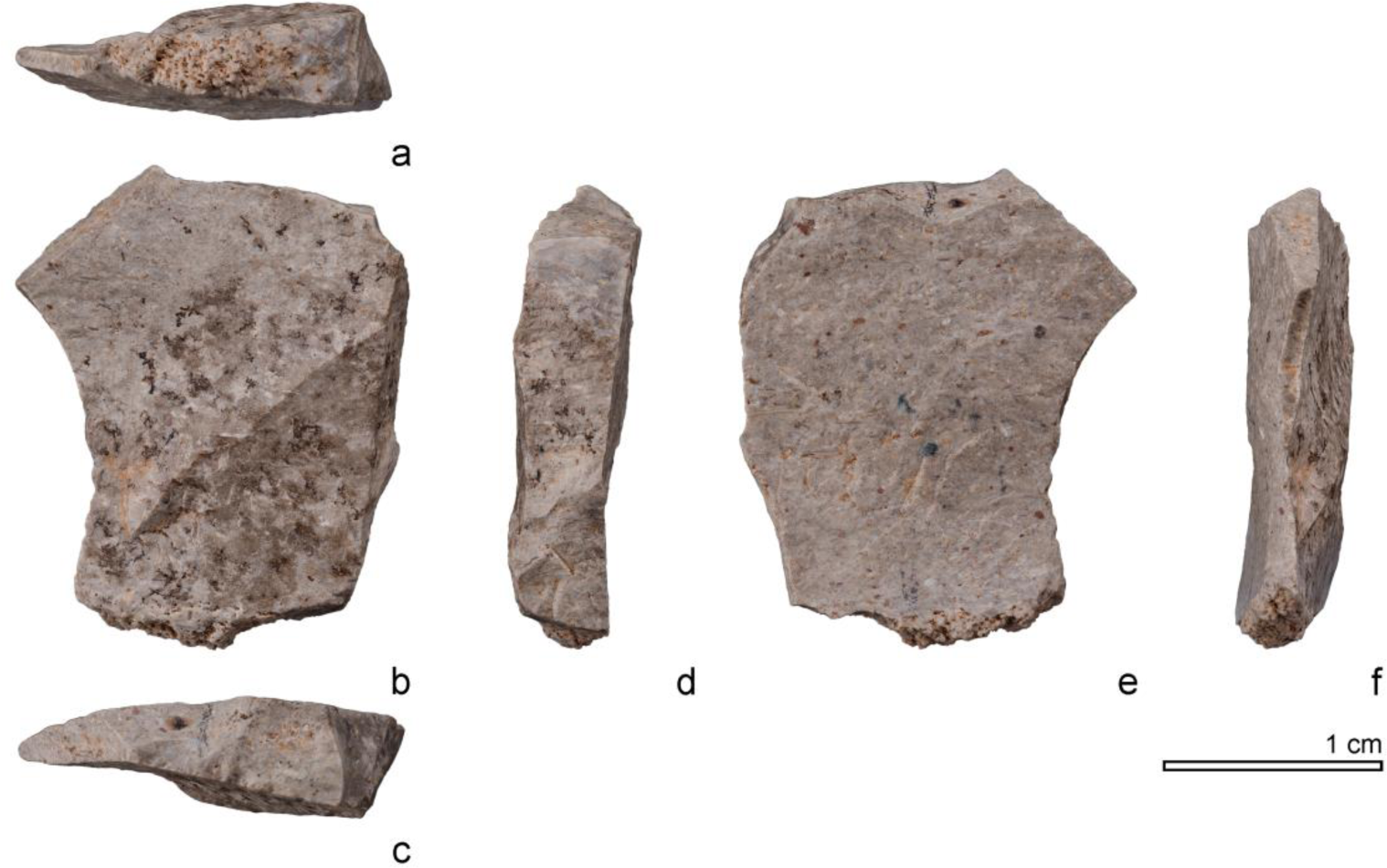
Limestone fragment, with no bulbus, no platform, no retouch and with irregularly crushed edges, described by Garcia Garriga et al. ([30], 81) as displaying “regular and continuous retouches at a steep angle” (their Fig 7e).

**Figure 12.**
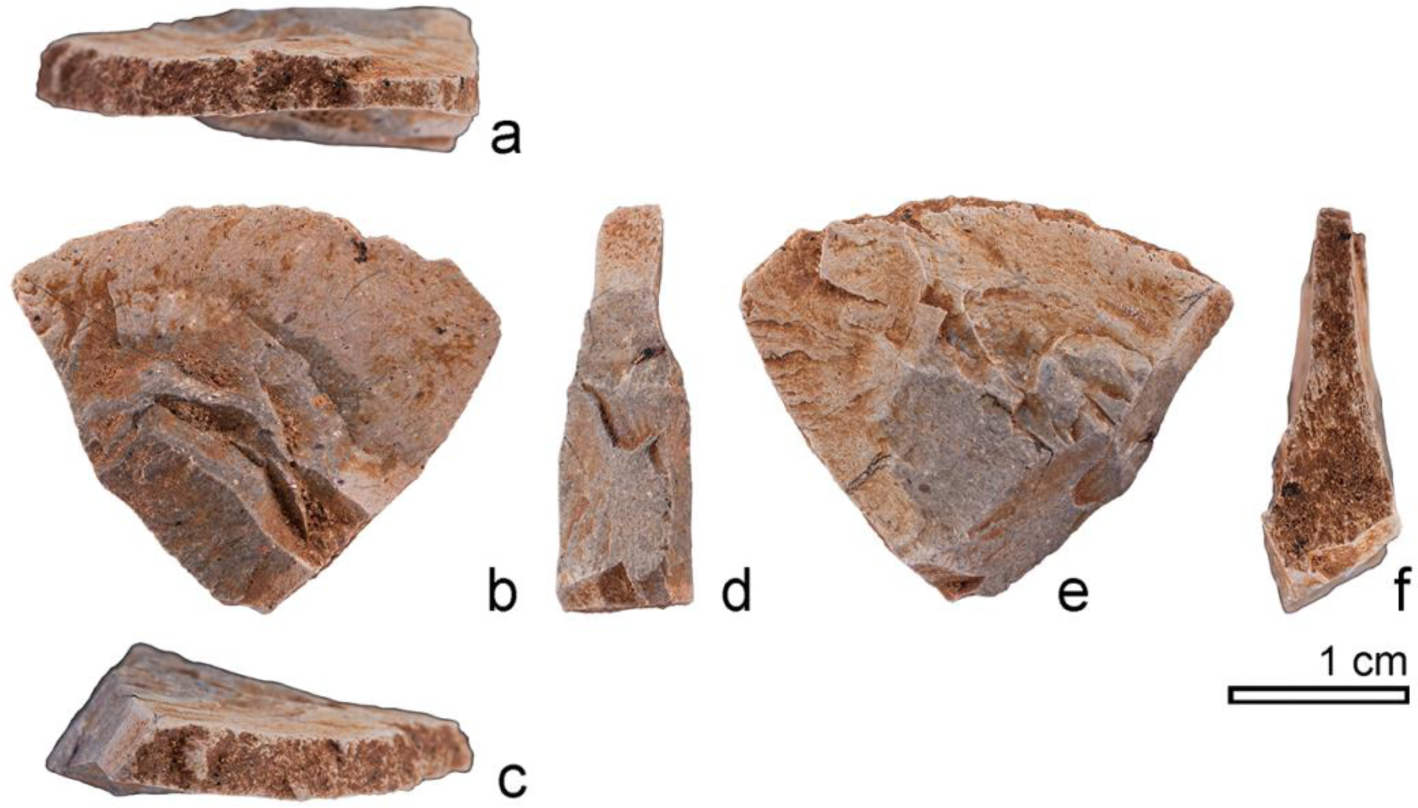
Chert fragment, interpreted as a flake produced in a “bipolar on an anvil technique” (compare Fig 6b in [30]).

**Figure 13.**
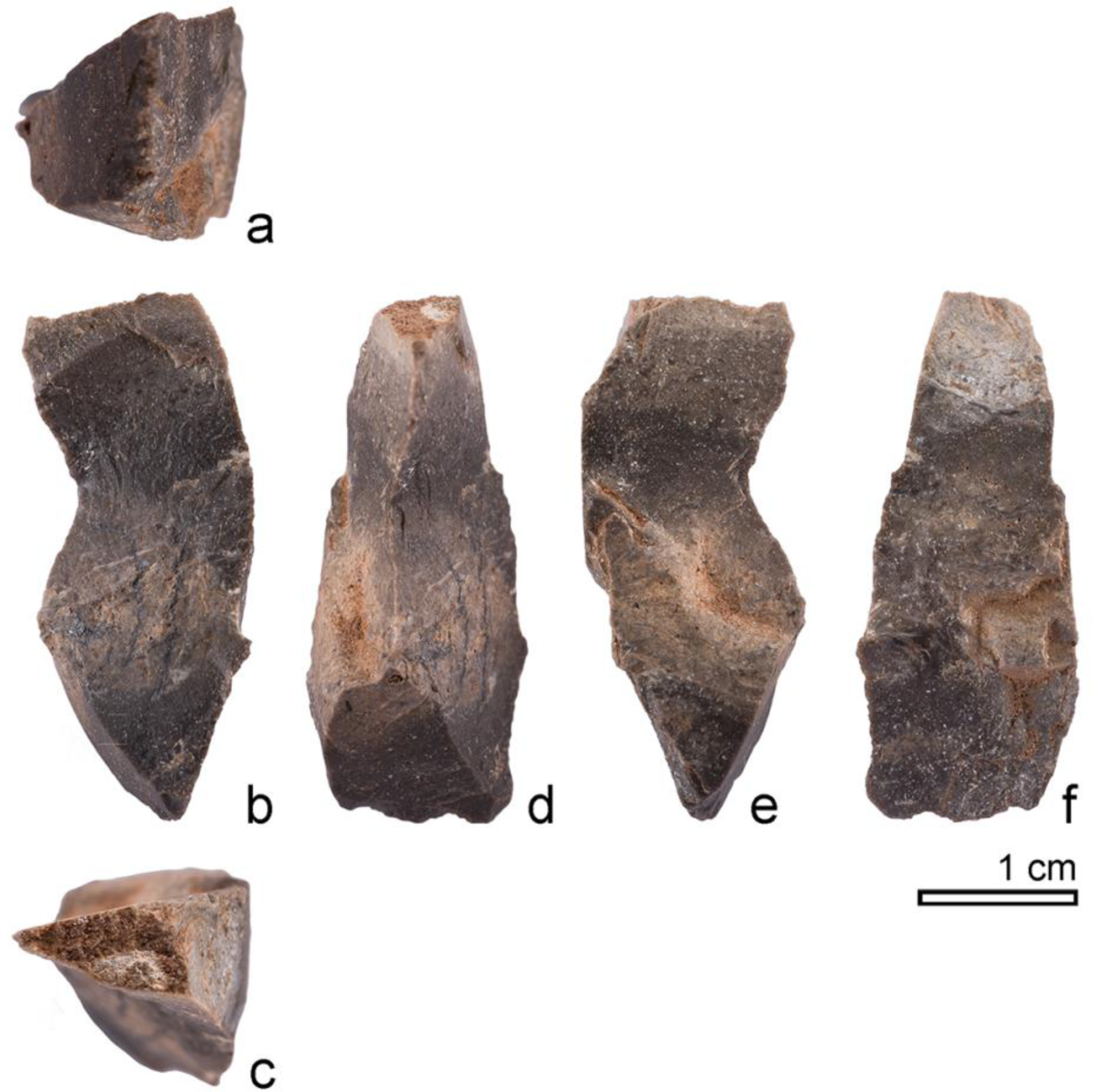
Chert fragment, interpreted as a flake produced in a “bipolar on an anvil technique” (compare Fig 6d in [30]).

**Figure 14.**
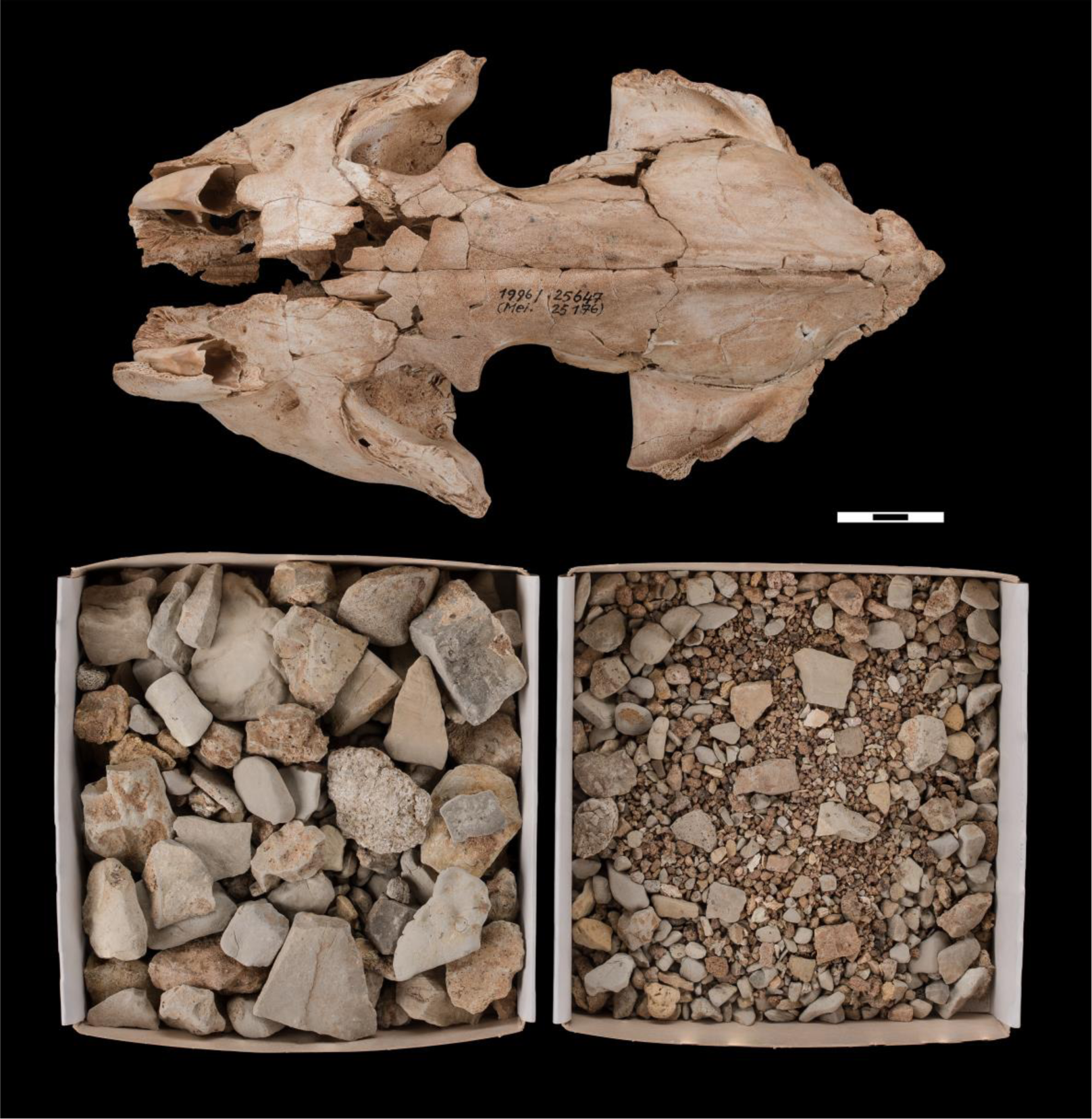
Untermassfeld, boxes with debris of Triassic limestone and chert, collected during preparation of a sediment block (50 x 35 x 25 cm) containing the fragmentary skull of a juvenile hyaena (*Pachycrocuta brevirostris*), excavated 21 July 1993 from squares 921 and 922, 0.40-0.50 cm above 0-level of the site (for exact find position see Kahlke 2006, foldout XIII), inventory number IQW 1996/25 647 (Mei. 25 176) (scale 3 cm).

## Discussion

### The Untermassfeld case

In summary, the studies claiming an early hominin presence at Untermassfeld are severely flawed in terms of data on provenance of the materials said to have been studied and in terms of (absence of) information on where the material is deposited. At least one of the faunal remains does come from the site, but the provenance of the lithics is completely unknown. The sample of faunal remains and lithics that we were able to study does not show any traces of hominin interference, and does not testify to a hominin presence at the site: we have no idea where the rest of the assemblage allegedly studied by Garcia Garriga and colleagues is stored and hence what it looks like, but based on the published finds that we were able to evaluate, Untermassfeld is not an archaeological site. As mentioned above, the Untermassfeld project has from the very beginning taken into consideration a possible presence of traces of hominin activities [42, 43], but more than three decades of fieldwork at the site, with 90 months of excavations there, as well as subsequent laboratory analyses by a wide range of specialists, so far did not yield any indication of a hominin presence in the fossil bearing deposits, not in terms of lithic artefacts, nor in hominin modifications of faunal remains. To clean up the record of the Early Pleistocene occupation of Europe, Untermassfeld should not be considered an archaeological site.

### Beyond Untermassfeld

Now more than two decades ago, the so-called Short Chronology [58] pushed for a critical evaluation of the record for the earliest occupation of Europe, in a large-scale evaluation of both dating evidence as well as evidence for the human workmanship of the lithic industries from the earliest sites [59]. The Short Chronology also questioned the artificial character of lithics from inferred early archaeological sites such as Kärlich (Germany) [60, 61], Prezletice (Czechia) [62] or Le Vallonnet in France [63], and suggested that these assemblages displayed all the characteristics of a selection of “primitive” pieces picked out from a matrix rich in rocks and pebbles, without any clear evidence for human agency at the sites – an issue that has been at stake in Palaeolithic archaeology since the days of the “eolith” controversies in the first decades of the 20^th^ century, whose main lesson was elegantly summarized by Warren [18], in a quote which is still highly topical almost a century later:

> “What is important, however, is the fact that such phenomena as the flaking of flints and occasional bulbs and also edge-knapping are produced by causes entirely apart from direct human effort. The likeness between the flaking produced by Nature and that produced by human agencies is sufficient to shift any burden of proof upon those who maintain the human origin of the stones; and this must be done not by a careful selection of picked specimens, but by a survey of the whole group” ([18],250).

The main conclusion of the Short Chronology was that most, if not all, the earliest published sites were either poorly dated or had lithics that were not of human manufacture, and that the oldest well-dated sites with artefacts were from around 500 ka or younger, suggesting a shift in the dynamics of occupation of this part of the Old World from around half a million years ago onward. The work set a rigorous standard against which new sites could be tested. The hypothesis however was soon falsified by new evidence from the Atapuerca TD sequence [64], with finds older than 800 ka, that passed the test and led to modification of the original model [65]. For northern Europe the threshold of 500-600 ka still held firm until new sites were discovered in the UK with evidence from Pakefield at ~600-700 ka [10, 66] and even older from Happisburgh Site 3 [9, 67]. It is important to underline that the hypothesis was falsified by new data from new sites as a result of new fieldwork, in the case of the Atapuerca TD sequence even explicitly aimed at falsifying the hypothesis [68], and not by new data from contested early sites, such as the ones mentioned above.

Solid datum points (sites) are important, as depending on the selection of “acceptable” sites, different scenarios emerge. Published scenarios include claims for a very early occupation of Europe, at about 1,4-1,5 Ma, on the basis of lithic materials from a few sites in southern Europe, such as Pirro Nord in southern Italy [69, 70] and lithic assemblages from exposures in the Guadix-Baza area near Orce in southern Spain [71, 72]. These finds would imply a hominin presence in Europe close in time to the earliest traces of a hominin presence in western Asia, at Dmanisi at the gates of Europe [73], but it needs to be stressed that the dating methods on which such claims are based are associated with large uncertainties [74, 75]. Well-dated unambiguous archaeological sites are significantly younger, suggestive of a significant (> 700,000 years) time lag between the Dmanisi finds and a post-Jaramillo first appearance of hominins in Europe, possibly prompted by profound environmental changes resulting from the high amplitude glacial oscillations of the late Early Pleistocene, around 900 ka [74]. Also on a smaller spatiotemporal scale, within Europe itself, tracking the presence and absence of hominins through time and in specific areas is an interesting exercise, as such differential occupation patterns are potentially very informative on hominin environmental limits and preferences [12, 19].

Fuelled by these issues, the database for early Europe is under constant maintenance through the normal process of scientific debate, which from time to time eliminates incorrect data, e.g. regarding the inferred arteficial character of finds (see for example the site Konczyce Wielkie 4 in South Poland [19]) or their age [76] [77], as well illustrated by the significant rejuvenation of the Ceprano (Italy) calvarium over the last decade by about 400,000 years, from an inferred pre-Brunhes-Matuyama [78] age to the current age estimate of around 400 ka [79] - see also the case of the Isernia site in Italy, which has moved from an inferred pre-Brunhes-Matuyama age [58] to well into the Middle Pleistocene, as recently demonstrated again by 40Ar/39Ar dating of the site [80].

The site of Untermassfeld has often been mentioned in one line with Vallparadís (Catalonia, Spain), as dating to the same time period and sharing a comparable “Mode 1” stone industry, “…which allows us to hypothesise that human groups with the same technology and acting as predators spread throughout the continent from the Jaramillo subchron onwards” ([30], 73). As at Untermassfeld, the primitive lithic assemblage from Vallparadis has been selected from stone rich (alluvial fan) deposits [32, 81], and as we do in this paper, the excavators of the Vallparadis site have stressed that the morphology of the lithics and the sedimentological data from the Vallparadis site do not support an anthropogenic origin of the recovered lithic pieces: they are not in primary context, fit very well in the general background of the lithic source area, and generally display rounded edges that cannot be accounted for by (the inferred) transport during a mud-flow event [82].

Issues regarding provenance, (non-) artefactual character and age of the “Oldowan” lithic assemblage from Vallparadis and other inferred early “archaeological sites” have been debated repeatedly in the last years [32, 74, 75, 81, 82], with Muttoni and colleagues recently concluding that currently “…there is no compelling chronological evidence of hominin presence before or during the Jaramillo” in Europe ([74],188).

The quality of the sites we put on our distribution maps counts, and hence database hygiene is crucial. The examples given above illustrate that there does exist an open debate that contributes to database maintenance in our field. In some prominent other cases one can observe a persistent absence of debate though. Once sites are in the scientific literature, claims can have very long half-lives. The French cave site of Le Vallonnet [83] is still a very prominent one in studies of early Europe, routinely treated as an archaeological site from the Jaramillo subchron, even though the palaeomagnetic dating evidence is debated [75, 77]. More importantly, the artificial character of the small and primitive (but well-published) lithic assemblage [83] from the site is ambiguous [58]. The lithics were already addressed in the Short Chronology and have also been critically reviewed in another publication [84] which has found no response from the Le Vallonet team yet:

> “The evidence for human activity at La Grotte du Vallonet would appear to be based on selection of modified lithic objects, that when examined, demonstrate a remarkable similarity to those from an overlying layer of Miocene conglomerate or ‘puddingstone’ containing pebbles with natural fractures, that were unmistakably produced millions of years before the first Australopithecines” [84] (p.76).

Beside issues regarding the artificial character of the lithics from Le Vallonet [58, 84], it has also been pointed out that the identification of an inferred bone tool from the site [83] was not based on taphonomic and contextual analyses of the bone assemblage and that the morphology of the cave (cave ceiling was less than 1 m above the layer in which the bone tool was found) is not easy to reconcile with a view of the site as a human occupation site [85]. The fact that these publications have never been mentioned in papers by the excavators of Le Vallonet, more than 20 years on, illustrates that there is still ground to gain for an open scientific debate on the earliest occupation of Europe.

### Conclusion

The absence of any hominin traces whatsoever at the Untermassfeld site and the problematic character of the claims made for sites such as Vallparadis and Le Vallonet imply that at the scale of Europe, solid evidence for a hominin presence in the Early Pleistocene is indeed rare [74], suggestive of an intermittent presence, with the earliest sites located at most 40 degree North - as is the case across much of Eurasia, from northern Spain (Atapuerca) [86] , to Dmanisi in the Georgia [73] all the way to the Nihewan Basin in northern China [87]. In the very final part of the Early Pleistocene hominins, at around 800-900 ka, may have expanded their range temporarily northward, following the European coastal areas, when conditions permitted. It is only much later, around 600 ka, that the record changes significantly, with an increase in site numbers all over western Europe, suggestive of changes in the character of hominin presence in this part of the world. These archaeological changes occur around the time period of the emergence of the Neanderthal lineage, which can be seen as independent – palaeontological- evidence for continuity of hominin occupation from that time period onward, minimally at the scale of Europe. Neanderthal populations expanded their range eastward, into the central parts of Europe from the middle part of the Middle Pleistocene, ~350 ka, onward, incorporating more challenging continental environments, [6, 12, 24], an expansion that has been related to the development of new cultural and possibly biological adaptations [13, 25–27].

We need solid data – well-dated sites with unambiguous traces of a hominin presence - to test the strength of the pattern described above, in fact a working hypothesis continuously subjected to testing and refinement by new field- and laboratory studies. In a discipline with limited access to original finds and especially their original context, scientists are heavily depending on the quality of scientific publications and hence on solid peer review. The case of Untermassfeld presented in extenso here underlines that peer review is an imperfect shield and that flawed claims do make their way into the scientific literature. Once such claims are there it is difficult to withdraw them from the scientific discourse, especially when published in high impact journals. The Untermassfeld case is admittedly an extreme one, but it does underline the importance of very careful peer review and the necessity of detailed description of the provenance of published materials as well where they are deposited for other researchers to access (see also: [88]).

A more general question is how to deal with disputed sites in the literature on earliest Europe. The claims for contested sites like Prezletice or Le Vallonet, for instance, have not become more valid because of the rich and well-documented finds from contemporaneous late Early and early Middle Pleistocene deposits in the Atapuerca site complex in Spain. This is not only an issue of the chronology of early occupation, but also about establishing the character of early occupation of a given area – continuous or discontinuous [3] - and the environmental tolerances and preferences of the early occupants of Eurasia. Chronology is crucial if we want to establish what triggered colonization of the various areas of Eurasia, under which circumstances various areas became populated and subsequently depopulated after earlier colonization, and whether spatiotemporal patterns of occupation correlate with new biological and /or cultural adaptations.

An open scientific debate is the most powerful tool to clean up and maintain our database, and that sometimes involves presenting negative evidence, as we have done here: *Untermassfeld is not an archaeological site*. Which site is next?

## Acknowledgements

The authors are grateful to N. Bicho (Faro), N. Conard (Tübingen), R. Klein (Stanford) and E. Pop (Monrepos/Leiden) as well as to the PLoS One reviewers (including M. Mussi, Rome) for commenting upon an earlier draft of the paper. R.-D.K. thanks J.-A. Keiler (Weimar) for the compilation of data on the thefts at the Untermassfeld excavation site, as well as for years of cooperation with the responsible police authorities and the Thuringian State Office for Monument Preservation and Archaeology. Further thanks go to S. Döring, E. Haase and T. Korn (all Weimar), to E. Noack and N. Viehöver (both Monrepos) as well as to J. Porck (Leiden) for producing the figures.

An earlier version of this paper was submitted as a News and Views article to the *Journal of Human Evolution* as a reaction on the Landeck and Garcia 2016 paper published in that journal [28] The *Journal of Human Evolution* Editors decided not to send our paper out for review, as we refused to follow their request to strike a section dealing with the problematic provenance history of the assemblage studied by Garcia Garriga and Landeck. We consider that provenance history to be an important part of the context of the study by Garcia and Landeck, including that the whereabouts of almost all the faunal material published in the *Journal of Human Evolution* is unknown, not deposited in a museum or other repository, and not accessible for other researchers.

